# A modified Arrestin1 Slows Retinal Degeneration Across Genetically Distinct Mouse Models of Inherited Retinal Disease

**DOI:** 10.64898/2026.11.10.765647

**Authors:** Ruining Peng, Kristin Ates-Hicks, Seok Hong Min, Stephanie Wohlgemuth, Astra Dinculescu, W. Clay Smith

## Abstract

Inherited retinal diseases (IRDs) are a genetically heterogeneous group of disorders caused by mutations in more than 340 genes, limiting the applicability of gene-targeted therapies and creating the need for broader treatment strategies. Photoreceptor dysfunction is a hallmark of IRDs and often results in disrupted retinal metabolic homeostasis, as photoreceptors have exceptionally high metabolic demands and provide metabolic resources to surrounding retinal cells. Consequently, enhancing retinal metabolism has emerged as a promising gene-agnostic strategy for treating IRDs. We previously developed a modified arrestin1 (ArrGG) that relieves arrestin1-mediated inhibition of the glycolytic enzyme enolase1, thereby increasing glycolytic output and slowing retinal degeneration in the *Rho^P23H/+^*mouse model. Here, we investigated the therapeutic potential of ArrGG as a gene-agnostic approach targeting endogenous metabolic enzyme activity across genetically distinct models of IRD.

Using two complementary photoreceptor-enriched transcriptomic analyses, we found no widespread, reproducible transcriptional changes following ArrGG treatment. These findings support our hypothesis that ArrGG modulates photoreceptor metabolism without inducing substantial transcriptional remodeling. The therapeutic efficacy of ArrGG was evaluated in three IRD mouse models representing rapid rod degeneration (rd10), cone degeneration (GC1KO), and RPE-associated retinal degeneration (rd6). ArrGG treatment was associated with preserved retinal function and slowed structural degeneration across all three models. Together, these findings support the potential of modulating endogenous metabolic enzyme activity to preserve photoreceptors across genetically distinct IRD models. This work suggests that ArrGG could be a promising gene-agnostic metabolic therapy for slowing retinal degeneration and preserving visual function in inherited retinal diseases.

## Introduction

Preserving photoreceptors through durable therapeutic strategies remains a major challenge in the treatment of inherited retinal diseases (IRDs). IRDs comprise a heterogeneous group of inherited visual disorders characterized by progressive photoreceptor degeneration and irreversible vision loss^1^, affecting approximately 1 in 2,000 individuals worldwide^2, 3^. Recent advances in gene-targeted therapy, such as the first FDA-approved treatment for RPE65-associated Leber congenital amaurosis using voretigene neparvovec (Luxturna^TM^), have demonstrated the clinical potential of mutation-specific therapies^4, 5^. However, more than 340 disease-causing genes have been identified to date (RetNet, https://retnet.org/disease), highlighting the remarkable genetic and clinical heterogeneity of IRDs. This genetic diversity poses a major challenge for gene-specific therapeutic strategies and motivates the development of more broadly applicable therapeutic interventions^6^.

One promising approach to overcome this challenge, regardless of the underlying genetic mutation, is to target retinal metabolism^6, 7^. The retina is one of the most energetically demanding tissues in the body^8^, and photoreceptors rely heavily on aerobic glycolysis^9^, making them especially susceptible to metabolic disturbances. Increasing evidence suggests that metabolic dysfunction is a common downstream consequence of genetically diverse IRDs, disrupting the metabolic homeostasis required for photoreceptor survival and contributing to progressive retinal degeneration^10, 11^. Studies have demonstrated that enhancing retinal metabolism through modulation of the insulin/mTOR pathway^12, 13^ or increasing glucose uptake through GLUT1 overexpression^14^ can preserve photoreceptors and improve retinal function in experimental models of IRD. However, most previously investigated approaches enhance retinal metabolism indirectly through modulation of signaling pathways or gene expression, leaving direct modulation of endogenous metabolic enzyme activity in photoreceptors largely unexplored.

We previously identified visual arrestin1 as an endogenous inhibitor of the glycolytic enzyme enolase1, reducing its enzymatic activity by approximately 25% in photoreceptors^15^. Structural analyses further demonstrated that enolase1 interacts with a surface of arrestin1 opposite its binding interface with photoactivated rhodopsin, providing a structural rationale for modifying the enolase1 interaction without necessarily disrupting canonical rhodopsin signaling^16^. Two amino acid residues, Glu-361 and Asp-362, contribute to steric interference with substrate access to the active site of enolase1^16^. Substituting these two residues with glycine (ArrGG) relieved the inhibitory effect on enolase1 catalysis while retaining enolase1 and rhodopsin binding^16^. Subsequent studies demonstrated enhanced glycolytic activity and slower retinal degeneration following ArrGG expression in the *Rho^P23H/+^* mouse model of IRD^17^.

Based on our previous findings, we hypothesized that direct modulation of endogenous metabolic enzyme activity to enhance photoreceptor glycolysis would not induce widespread changes in the photoreceptor transcriptome. Furthermore, enhancing glycolysis through ArrGG expression could improve photoreceptor energy metabolism while increasing lactate availability to support neighboring retinal cells. Because ArrGG targets enolase1 independently of the disease-causing mutation, we proposed that ArrGG treatment could slow photoreceptor loss and retinal degeneration across genetically diverse forms of IRD, including diseases characterized by primary rod degeneration, cone degeneration, or retinal defects originating outside photoreceptors.

In this study, we investigated whether ArrGG expression induced widespread changes in the photoreceptor transcriptome and evaluated its potential to preserve retinal function and slow photoreceptor degeneration across three genetically distinct mouse models of IRD.

## Methods

### Animals

This study was conducted with approval from the University of Florida Institutional Animal Care and Use Committee (IACUC) and in accordance with the Association for Research in Vision and Ophthalmology (ARVO) Statement for the Use of Animals in Ophthalmic and Vision Research.

### Mouse Strains

C57BL/6J mice were obtained from the internal breeding colony maintained by Animal Care Services (ACS) at the University of Florida. Rod-specific RiboTag mice were generated by crossing RiboTag mice [B6J.129(Cg)-*Rpl22^tm^*^1^*^.1Psam^*/SjJ; The Jackson Laboratory, Stock No. 029977] with rhodopsin-iCre mice [B6.Cg-*Pde6b^+^*Tg(Rho-icre)1Ck/Boc; The Jackson Laboratory, Stock No. 015850]. The resulting mice expressed hemagglutinin (HA)-tagged Rpl22 predominantly in rod photoreceptors. The genotype was confirmed by polymerase chain reaction (PCR), as previously described^18^, and the mice were maintained as an in-house breeding colony. The rd6 [B6.Cg-*Mfrp^rd^*^6^/J; The Jackson Laboratory, Stock No. 003684] and rd10 [B6.CXB1-*Pde6b^rd^*^10^/J; The Jackson Laboratory, Stock No. 004297] mouse strains were maintained as in-house breeding colonies at the University of Florida. The rd10 mice were housed under dim light conditions. GC1KO mice were kindly provided by Dr. Shannon E. Boye at the University of Florida.

### Generation of a Stable ARPE-19 Cell Line Expressing ArrWT

ARPE-19 cells were maintained in BenchStable™ Dulbecco’s Modified Eagle Medium/Nutrient Mixture F-12 (DMEM/F12; Gibco, Cat. No. A4192001) supplemented with 10% fetal bovine serum (FBS; Gibco, Cat. No. A5670701) and 1× Antibiotic-Antimycotic (Gibco, Cat. No. 15240-062) at 37°C in a humidified atmosphere containing 5% CO₂. Cells were transduced at a multiplicity of infection (MOI) of 10 with a lentiviral vector encoding FLAG-tagged wild-type arrestin1 (ArrWT) under the control of the chicken β-actin (CBA) promoter, with a 2A-linked puromycin resistance gene. The lentiviral vector was custom-designed, constructed, and packaged by VectorBuilder (Chicago, IL, USA). Following transduction, cells were selected with 5 μg/mL puromycin (Gibco, Cat. No. A11138-03). Puromycin-resistant cells were expanded and subsequently maintained in culture medium containing 1 μg/mL puromycin. Stable expression of FLAG-tagged ArrWT was confirmed by Western blotting using an anti-FLAG antibody (0.1 μg/mL; Proteintech, Cat. No. 66008-4-Ig), with an anti-GAPDH antibody (Proteintech, Cat. No. 10494-1-AP) as a loading control.

### Lactate Secretion Assay

Parental ARPE-19 cells and stable ArrWT-expressing ARPE-19 cells were seeded in 12-well plates (Corning, Cat. No. 3515) at a density of 2 × 10^5^ cells per well. After 24 h, a subset of the ArrWT-expressing ARPE-19 cells was transiently transfected with an ArrGG-expressing plasmid driven by the CBA promoter using Lipofectamine^TM^ 3000 Transfection Reagent (Invitrogen, Cat. No. L3000015). Twenty-four hours after transfection, the culture medium was replaced with fresh medium containing 5 μM rotenone (Sigma-Aldrich, Cat. No. R8875) to inhibit mitochondrial electron transport. Conditioned medium was collected at 30, 45, 60, 75, 90, 105, and 120 min after medium replacement. Lactate concentrations in the collected medium were determined using the Lactate-Glo™ Assay (Promega, Cat. No. J5021) according to the manufacturer’s instructions. Lactate secretion rates were calculated by linear regression of extracellular lactate concentration against collection time, with the slope representing the lactate secretion rate. The calculated rates were normalized to those of parental ARPE-19 cells. Statistical comparisons among experimental groups were performed using one-way analysis of variance (ANOVA) in GraphPad Prism (v10.3.1).

### Seahorse XF Glycolytic Rate Assay

Parental ARPE-19 cells, ArrWT-expressing ARPE-19 cells, and ArrWT-expressing ARPE-19 cells transiently transfected with ArrGG were analyzed using the Seahorse XF Glycolytic Rate Assay Kit (Agilent Technologies, Cat. No. 103344-100). Cells were seeded in Seahorse XFe96/XF Pro cell culture microplates (Agilent Technologies, Cat. No. 103794-100) at a density of 2.5 × 10^4^ cells per well 24 h before the assay. On the day of analysis, the culture medium was replaced with Seahorse XF assay medium supplemented with 1 mM pyruvate, 2 mM glutamine, and 10 mM glucose, and cells were incubated at 37°C in a non-CO_2_ incubator for 1 h before measurement.

Extracellular acidification rate (ECAR) and oxygen consumption rate (OCR) were measured in real time to determine the glycolytic proton efflux rate (glycoPER). Basal glycolysis was measured before mitochondrial inhibition, and compensatory glycolysis was determined following the injection of rotenone and antimycin A. A subsequent injection of 2-deoxy-D-glucose (2-DG) was used to inhibit glycolysis and account for non-glycolytic acidification. Glycolytic measurements were normalized to the number of viable cells determined after completion of the assay. The compensatory glycolytic response was calculated as the percentage increase in compensatory glycolysis relative to basal glycolysis. Values were further normalized to those of parental ARPE-19 cells and compared among experimental groups using one-way ANOVA in GraphPad Prism (v10.3.1).

### Construction of AAV vectors

AAV2(quad+T-V) vectors expressing ArrWT or ArrGG under the control of the human opsin promoter were constructed and produced as previously described^17, 19^. The AAV-GFP vector was generated using the same vector backbone, with the arrestin1 coding sequence replaced by that of green fluorescent protein (GFP) coding sequence. All AAV vectors were packaged, purified, processed to remove endotoxins, and titrated by the Ocular Gene Therapy Core at the University of Florida as previously described^20^.

### Intravitreal Injections

Intravitreal injections were performed as previously described by Nelson et al.^17^. Briefly, mouse eyes were dilated with 0.25% phenylephrine HCl and 1% tropicamide before anesthesia and again immediately before injection. Mice were anesthetized by intraperitoneal injection of xylazine (5–10 mg/kg) and ketamine (70–100 mg/kg) prepared in sterile saline solution. Under an operating microscope, the needle was passed through the sclera posterior to the limbus and advanced into the vitreous cavity. A total volume of 1 μL of AAV vector (ArrWT: 7.51 × 10^9^ vg/eye; ArrGG: 2.87 × 10^10^ vg/eye; GFP: 1.03 × 10^8^ vg/eye) or 1 μL of balanced salt solution (BSS) was injected into the vitreous cavity. GenTeal gel (0.3%; Novartis) was applied to the corneal surface to facilitate visualization during injection and prevent corneal drying.

### Photoreceptor Enrichment by FACS

C57BL/6J mice received intravitreal co-injections of AAV-ArrGG and AAV-GFP mixed at a 1:1 volume ratio, with AAV-GFP serving as a fluorescent marker to facilitate the identification and enrichment of AAV-transduced photoreceptors. Control mice received AAV-GFP alone. Two weeks after injection, retinas were dissociated into single-cell suspensions using papain (Worthington, Cat. No. LK003150)^21^ according to the manufacturer’s instructions. Three freshly isolated retinas were pooled per sample, washed with ice-cold phosphate-buffered saline (PBS), and incubated in pre-equilibrated papain solution at 37°C for 20 min with gentle mixing every 5 min. Following enzymatic digestion, retinal tissue was gently triturated, and papain activity was quenched with ovomucoid-containing Earle’s Balanced Salt Solution (EBSS). The resulting cell suspension was passed through a 40-μm cell strainer to remove undissociated tissue and cellular aggregates.

Dissociated retinal cells were collected by centrifugation at 300 × *g* for 4 min and resuspended in 1% bovine serum albumin (BSA). Cells were incubated with an anti-CD73 antibody (BioLegend, Cat. No. 127208) for 15 min on ice and protected from light. Cells positive for both CD73 and GFP were sorted to enrich for AAV-transduced photoreceptors using a Sony SH800 cell sorter (Sony Biotechnology, CA, USA) at the University of Florida ICBR Cytometry & Optical Microscopy Core Facility (RRID: SCR_019119). Photoreceptor RNA was isolated using the PicoPure RNA Isolation Kit (Applied Biosystems™, Cat. No. KIT0204). ArrGG transgene expression in the sorted photoreceptors was confirmed by reverse transcription quantitative polymerase chain reaction (RT-qPCR) (sense primer: 5’-CATGCACCCTCAGCCTGGCGGCCCAGCTAAGGAAAGTTATC-3’; antisense primer: 5’-TCACAGATCTTCCTCAGAGATAAGTTTTTGCTCCTCATCAACGTCATTCTTGTCTCTC-3’).

### RNA Isolation from Rod-Specific RiboTag Mice

RiboTag immunoprecipitation was performed following the protocol described by Rajala et al.^18^. Briefly, rod-specific RiboTag mice received intravitreal injections of AAV-ArrGG, while control mice received BSS. Four weeks after injection, four retinas were pooled per sample and incubated in DMEM containing 100 μg/mL cycloheximide (Sigma-Aldrich, Cat. No. 239765) at 37°C for 10 min. Retinas were then collected by centrifugation at 300 × *g* for 2 min, flash-frozen in liquid nitrogen, and pulverized using a handheld homogenizer. Retinal tissue was resuspended in 200 μL of polysome buffer containing 50 mM Tris-HCl (pH 7.5), 100 mM KCl, 12 mM MgCl_2_, 1% Igepal CA-630, 1 mM dithiothreitol, 200 U/mL RNaseOUT, 1 mg/mL heparin sodium salt, 100 μg/mL cycloheximide, and an EDTA-free protease inhibitor cocktail prepared in DEPC-treated water. Lysates were then centrifuged at 15,000 rpm for 2 min at 4°C.

The cleared lysate was incubated with a purified mouse monoclonal anti-HA antibody (5 μL per 200 μL of lysate; GenScript, Cat. No. A01244) for 1 h at 4°C. Dynabeads^TM^ Protein G (Invitrogen, Cat. No. 10004D), pre-equilibrated in polysome buffer, were then added to the lysate and incubated for an additional 30 min at 4°C. The magnetic beads containing immunoprecipitated complexes were washed three times with high-salt buffer containing 50 mM Tris-HCl (pH 7.5), 300 mM KCl, 12 mM MgCl_2_, 1% Igepal CA-630, 1 mM dithiothreitol, 200 U/mL RNaseOUT, 1 mg/mL heparin sodium salt, 100 μg/mL cycloheximide, and an EDTA-free protease inhibitor cocktail. RNA was released by adding 500 μL TRI Reagent (Sigma-Aldrich, Cat. No. T9424) and purified using the PureLink^TM^ RNA Mini Kit (Invitrogen, Cat. No. 12183020) according to the manufacturer’s instructions. ArrGG expression in the rod photoreceptor-enriched, ribosome-associated RNA fraction was confirmed by RT-qPCR as described above.

### Photoreceptor RNA Sequencing

RNA sequencing was performed by the University of Florida ICBR Gene Expression & Genotyping Core Facility (RRID: SCR_019145) using the Illumina NovaSeq X Plus platform. Sequencing data were processed by the University of Florida ICBR Bioinformatics Core Facility (RRID: SCR_019120). Reads were processed using Cutadapt to remove sequencing adapters and low-quality bases with Phred quality scores below 20^22^. Reads shorter than 40 bases after trimming were excluded from further analysis. Cleaned reads were aligned to the *Mus musculus* reference genome (GRCm38/mm10) using STAR (v2.7.9a)^23^. Alignment results were subsequently processed using HTSeq (v0.11.2)^24^, SAMtools, and in-house scripts developed by the UF ICBR Bioinformatics Core to remove potential PCR duplicates and quantify uniquely mapped reads for gene expression analysis.

Three independent biological samples were analyzed per experimental group, with each sample comprising pooled retinas as described above. Downstream analyses were performed using R (v4.1.3). Principal component analysis (PCA) was used to evaluate sample clustering and detect potential outliers, and volcano plots were generated to visualize differences in gene expression between experimental groups. Differential gene expression analysis was performed using a DESeq2-based R pipeline^25^. Genes with |log₂FC| ≥ 1 and adjusted *P* ≤ 0.05 were considered differentially expressed.

### Quantitative Real-Time Reverse Transcription PCR Validation

Selected genes identified from the RNA-seq analysis were further evaluated by RT-qPCR using independent biological samples processed through the rod-specific RiboTag procedure described above (*n*=4). Total RNA was reverse-transcribed into complementary DNA (cDNA) using the First-Strand cDNA Synthesis Kit (Cytiva, Cat. No. 27926101) according to the manufacturer’s instructions. Quantitative PCR was performed using qPCRBIO SyGreen Blue Mix (Genesee Scientific, Cat. No. 17-505) on a CFX Real-Time PCR Detection System (Bio-Rad Laboratories, CA, USA). Primers used for amplification of the selected genes are listed in Table 1. Relative gene expression, 2^−ΔΔCq^, was calculated, with rhodopsin (*Rho*) used as the reference gene for normalization. All reactions were performed in duplicate, and the averaged Cq values of the two technical replicates were used for subsequent calculations. Relative expression values were compared between the AAV-ArrGG and control groups using multiple unpaired *t*-tests in GraphPad Prism (v10.3.1).

**Table 1.**
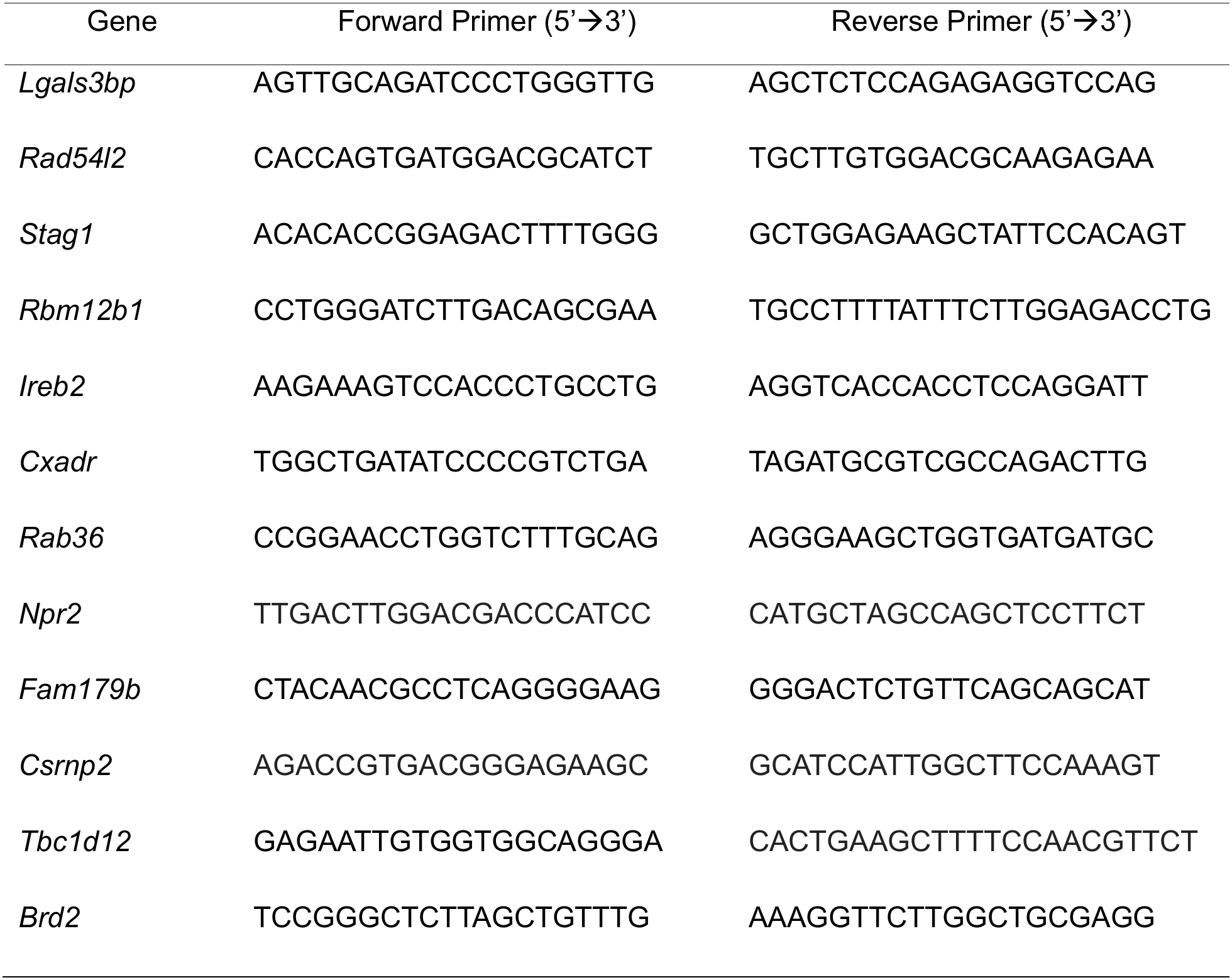
Primers for selected genes from photoreceptor RNA-seq analysis.

### Electroretinographic Analysis

Electroretinograms (ERG) were recorded using the Celeris ERG system (Diagnosys LLC, MA, USA) and Espion software (v6.68.2). Mice were dark-adapted overnight with food and water available. Before recording, pupils were dilated and mice were anesthetized as described above. Mice were placed on a 37°C heating platform to maintain body temperature throughout the procedure. Corneal electrodes were positioned in gentle contact with both eyes, and reference electrodes were placed subcutaneously at the head and tail.

Dark-adapted ERG responses were recorded at flash intensities of 0.003, 0.01, 0.03, 0.1, 0.3, 1, 3, 10, and 30 cd·s/m^2^. At the first six intensities, 10 flashes were delivered with a 5-s interstimulus interval. At the final three intensities, 15 flashes were delivered with a 30-s interstimulus interval. Following 8 min of light adaptation at 30 cd/m^2^, light-adapted ERG responses were recorded at 0.15, 0.5, 1.3, 3.5, 10, and 30 cd·s/m^2^ on a 30 cd/m^2^ background. Each intensity included 25 flashes with a 1-s interstimulus interval.

ERG a-wave amplitude was measured from the baseline to the negative trough, and b-wave amplitude was measured from the negative trough to the subsequent positive peak. Amplitudes of a-wave and b-wave were compared among groups using two-way ANOVA in GraphPad Prism (v10.3.1).

### Optical Coherence Tomography

Optical coherence tomography was performed using the HRA+OCT Spectralis system (Heidelberg Engineering GmbH, Heidelberg, Germany) at the indicated time points to measure retinal outer nuclear layer (ONL) thickness and evaluate retinal degeneration over time. Before imaging, pupils were dilated and mice were anesthetized as described above. ONL thickness was quantified at predefined distances from the optic nerve head along the temporal-nasal axis, as indicated for each experiment. Measurements at individual time points were analyzed using one-way ANOVA, and longitudinal measurements were analyzed using two-way ANOVA in GraphPad Prism (v10.3.1).

### Immunohistochemistry and Microscopy

Whole mouse eyes were dissected from the orbit and fixed in 4% paraformaldehyde (PFA) prepared in 2 × PBS (274 mM NaCl, 5.4 mM KCl, and 20 mM phosphate buffer, pH 7.4) at room temperature for 30 min. The cornea was then removed by making a circumferential incision along the limbus, and the resulting eyecups were returned to 4% PFA for an additional 3 h at 4°C. Following fixation, eyecups were embedded in O.C.T. compound (Fisher, Cat. No. 23730571) and cryosectioned at a thickness of 10 μm.

Retinal sections were stained with 1 μg/mL C10C10 mouse monoclonal anti-arrestin1 antibody^26^ or a 1:500 dilution of rabbit polyclonal anti-cone opsin antibody (Sigma-Aldrich, Cat. No. AB5405). Primary antibody binding was detected using 8 μg/mL Alexa Fluor^TM^488-conjugated goat anti-mouse IgG secondary antibody (Invitrogen, Cat. No. A11029) or 8 μg/mL Alexa Fluor^TM^488-conjugated donkey anti-rabbit IgG secondary antibody (Invitrogen, Cat. No. A21206). Nuclei were stained with 10 μg/mL DAPI (Invitrogen, Cat. No. D1306). Sections were imaged using a Leica SP8 confocal microscope (Leica Microsystems, Wetzlar, Germany), and Z-stacks were acquired at 0.5 μm intervals and displayed as maximum-intensity projections. Images from different experimental groups were acquired using identical laser power, detector gain, and other acquisition parameters, including scan resolution and averaging settings, to ensure consistency across groups.

### Retina Flat Mount Staining and Cone Quantification

Whole mouse eyes were enucleated and fixed in 4% PFA for 3 h, followed by a 30-min incubation in PBS. After fixation, the cornea and lens were removed, and the neural retina was carefully dissected from the eyecup. Retinal flat mounts were stained with 10 μg/mL biotinylated peanut agglutinin (PNA; Vector Laboratories, Cat. No. B-1075-5), followed by detection with 10 μg/mL Fluorescein Avidin D (Vector Laboratories, Cat. No. A-2001-5). Flat mounts were imaged with a Leica SP8 confocal microscope and a Leica widefield fluorescence microscope (Leica Microsystems, Wetzlar, Germany).

PNA-positive cone structures were manually quantified within three predefined 8.5 x 10^4^ μm^2^ regions from each retinal flat mount. Counts from the three regions were averaged to obtain a single value for each retina. Three independent retinas were analyzed per experimental group (*n*=3). Measurements were compared among treatment groups using one-way ANOVA in GraphPad Prism (v10.3.1).

### Optomotor Response Testing

Visually guided behavior was assessed by measuring optomotor responses using the OptoMotry system (CerebralMechanics, Lethbridge, Canada), following a previously described method^27^. Mice were placed on an elevated central platform and allowed to acclimate for 2 min before testing. Rotating vertical gratings were presented at 100% contrast, a rotation speed of 12°/s, and a previously reported mean stimulus luminance of 70 cd/m², with spatial frequencies ranging from 0.003 to 0.7 cycles per degree. The presence or absence of visually guided head-tracking responses to clockwise or counterclockwise rotation of the gratings was recorded. Spatial frequency thresholds were determined using a simple staircase procedure, and visual acuity was defined as the highest spatial frequency that elicited a consistent tracking response. Each mouse underwent three independent tests, and the resulting thresholds were averaged for statistical analysis. The investigator was blinded to treatment allocation to minimize observer bias. Mean visual acuity thresholds at individual time points were analyzed using one-way ANOVA and longitudinal measurements were analyzed using two-way ANOVA in GraphPad Prism (v10.3.1).

## Results

### Expression of ArrGG in the Presence of Wild-Type Arrestin1 in ARPE-19 Cells

We have previously demonstrated that ArrGG can be used to increase lactate production in the retina, which is consistent with the disinhibition of enolase1 leading to an increased glycolytic rate. To provide an empirical test of ArrGG’s enhancement of glycolysis, we elected to use a defined cell culture system in which we could manipulate arrestin1. We chose ARPE-19 cells since these cells do not normally express arrestin1. This cell model allowed us to compare glycolytic activity in parental cells, cells expressing arrestin1 (ArrWT) alone, and cells co-expressing ArrWT and ArrGG, thereby evaluating the effects of introducing ArrWT and subsequently ArrGG on glycolytic activity. To establish this model, we generated ARPE-19 cells stably expressing FLAG-tagged ArrWT using a lentiviral vector (**Fig. 1A**). Stable expression of ArrWT in ARPE-19 cells was confirmed by anti-FLAG immunoblotting (**Fig. 1B**, compare lanes 1-3 with lanes 4-6).

**Figure 1.**
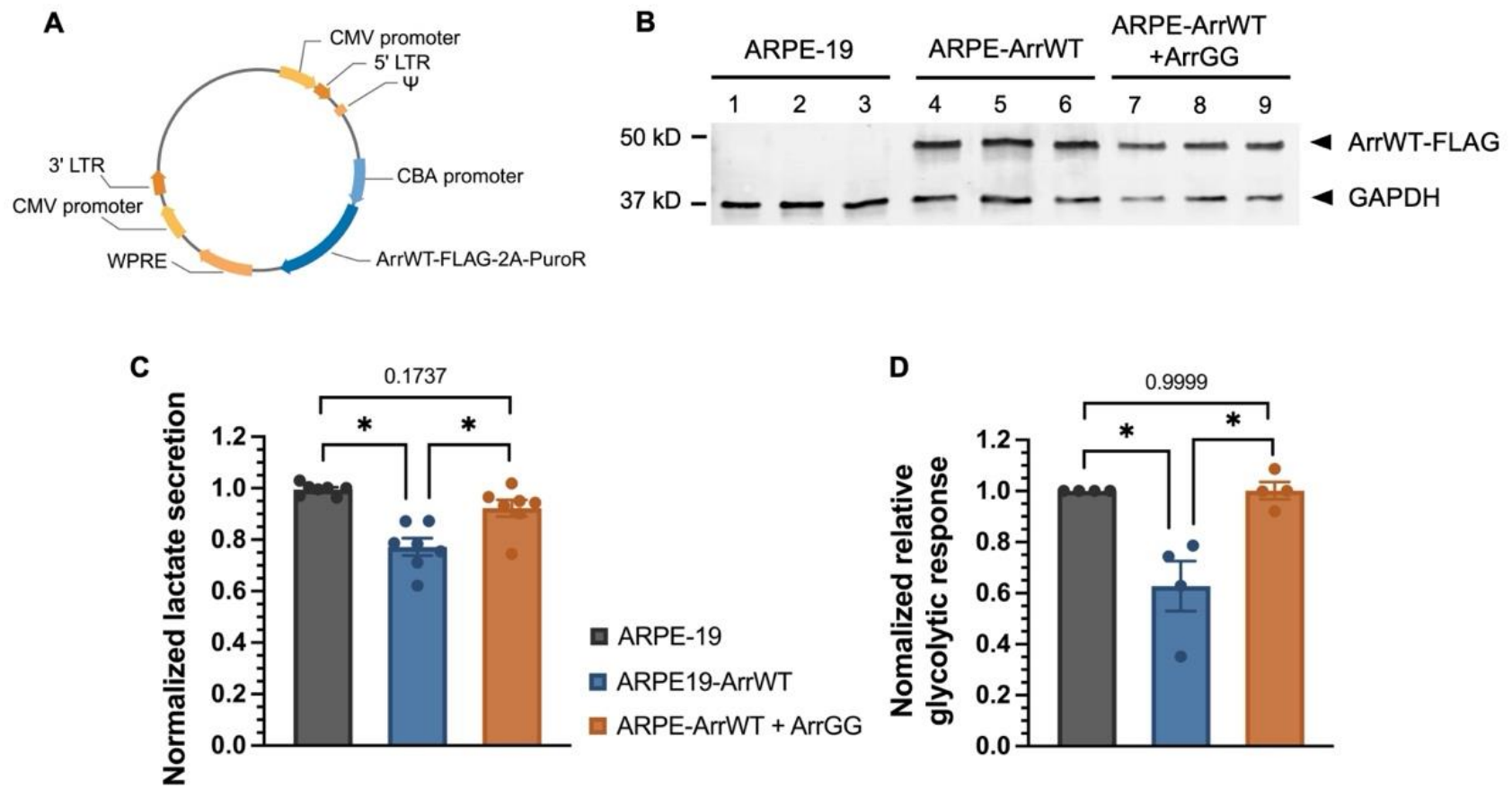
ArrGG maintains glycolytic responses in the presence of ArrWT. (**A**) Schematic of the transfer plasmid used to prepare lentivirus for stable expression of FLAG-tagged wild-type arrestin1 (ArrWT). The vector contains a CBA promoter driving the ArrWT-FLAG-2A-PuroR expression cassette together with elements required for lentiviral vector production and transgene expression. **(B)** Immunoblots of lysates from ARPE-19 cells (**lanes 1-3**), ARPE-19 cells stably expressing ArrWT (ARPE-ArrWT; **lanes 4-6**), and ARPE-ArrWT cells transfected with an ArrGG-expressing plasmid (ARPE-ArrWT + ArrGG; **lanes 7-9**). Blots were simultaneously probed with anti-FLAG and anti-GAPDH antibodies. **(C)** Rates of lactate secretion were measured in ARPE-19 cells, ARPE-ArrWT cells, and ARPE-ArrWT cells expressing ArrGG (*n*=7). Values were normalized to the ARPE-19 control. **(D)** Seahorse analysis of glycolytic responses derived from glycoPER measurements following mitochondrial inhibition in ARPE-19 cells, ARPE-ArrWT cells, and ARPE-ArrWT cells expressing ArrGG (*n*=4). Values were normalized to the ARPE-19 control. Data are presented as mean ± SEM. Statistical significance was determined by one-way ANOVA (*: *P*<0.05).

The cell model was subsequently used to evaluate whether ArrGG could increase glycolytic activity in the presence of ArrWT. First, we measured the rate of lactate secretion as an indicator of glycolytic output. Addition of ArrWT to ARPE-19 cells reduced the rate of lactate secretion by 22.4% compared with parental ARPE-19 cells (**Fig. 1C**, compare gray bar to blue bar). This value closely matches the ∼25% decrease in enolase1 catalytic activity previously observed in reconstitution assays using purified arrestin1 and enolase1 proteins^15^ and in HEK-293T cells transiently transfected with arrestin1^16^.

To determine whether the change in the rate of lactate secretion could be ascribed to an effect on glycolysis, we next performed an independent Seahorse Glycolytic Rate Assay to quantify the glycolytic response to mitochondrial inhibition^28^. Compared with parental ARPE-19 cells, cells stably expressing ArrWT showed a 37.2% reduction in the relative glycolytic response (**Fig. 1D**, compare gray bar to blue bar), indicating that the reduced rate of lactate secretion in cells expressing ArrWT is associated with reduced glycolytic activity.

To then determine if ArrGG could counteract the effect of ArrWT, we then transiently transfected the ArrWT-ARPE19 cells with ArrGG. Transient expression of ArrGG increased the rate of lactate secretion back to that of the ARPE-19 cells without arrestin1 (**Fig. 1C**, orange bar). Seahorse analysis of the ArrWT cells transfected with ArrGG showed a 59.5% increase in the glycolytic response to a level comparable to that of parental ARPE-19 cells (**Fig. 1D**, orange bar). These findings are consistent with the lactate secretion assay, independently supporting the conclusion that ArrGG restores the glycolytic response suppressed by ArrWT. Since it is possible that transient transfection with ArrGG could suppress ArrWT expression, we also assessed whether transient transfection with an ArrGG plasmid altered ArrWT expression (**Fig. 1B**, lanes 7-9). In these cells, ArrWT expression was maintained following ArrGG transfection, comparable to that in cells not transfected with ArrGG (lanes 4-6).

Together, these findings support that the effect of arrestin1 on lactate secretion is through suppression of glycolysis and that overexpression of ArrGG can restore full glycolytic activity. This finding, combined with our previous biochemical demonstration that ArrGG retains its ability to bind enolase1 while eliminating the inhibitory effect observed with ArrWT on enolase1 catalytic activity^16^, supports our proposed mechanism in which ArrGG competitively displaces ArrWT from enolase1 and thus increases glycolytic activity.

### Photoreceptor Transcriptomic Profiling Following ArrGG Treatment

Our hypothesis for the efficacy of ArrGG in slowing retinal degeneration that we saw in the *Rho^P23H/+^*mouse model is that the increased glycolytic output benefits both photoreceptors and the surrounding RPE and Müller glia. However, it is also possible that AAV-vectored ArrGG impacts an unanticipated signaling pathway. Additionally, metabolic changes are frequently accompanied by transcriptional remodeling^29, 30^, which could also lead to alternative pathway activation. To assess the potential contributions of other pathways, we performed transcriptomic analyses to investigate whether ArrGG expression is associated with substantial changes in photoreceptor gene expression by comparing photoreceptors from ArrGG-treated to control retinas. For this analysis, ArrGG was packaged in an AAV2(quad+T-V) vector^19^ under the control of the human opsin promoter to deliver this modified arrestin1 to the mouse retina and drive photoreceptor-specific expression (**Fig. 2A**). C57BL/6J mice were intravitreally co-injected with AAV-ArrGG and AAV-GFP to fluorescently label photoreceptors for FACS sorting, whereas control mice received AAV-GFP alone. Two weeks after injection, retinas were dissociated and GFP-positive/CD73-positive photoreceptors were isolated and enriched for bulk RNA sequencing. To confirm ArrGG expression in the enriched cells, a portion of the RNA was reverse transcribed and analyzed by qPCR for the ArrGG transgene. As expected, cells isolated from ArrGG/GFP-treated eyes showed enriched expression of the ArrGG transgene, whereas those isolated from eyes injected with AAV-GFP alone did not (**Fig. 2A**). Transcriptomic profiles were then compared between the ArrGG/GFP-treated and GFP-treated groups. RNA-seq analysis identified no statistically significant differences in gene expression between the ArrGG/GFP-treated and GFP-treated groups, as none of the genes met the predefined thresholds for differential expression (|log₂FC| ≥ 1 and adjusted *P* ≤ 0.05) (**Fig. 2C**).

**Figure 2.**
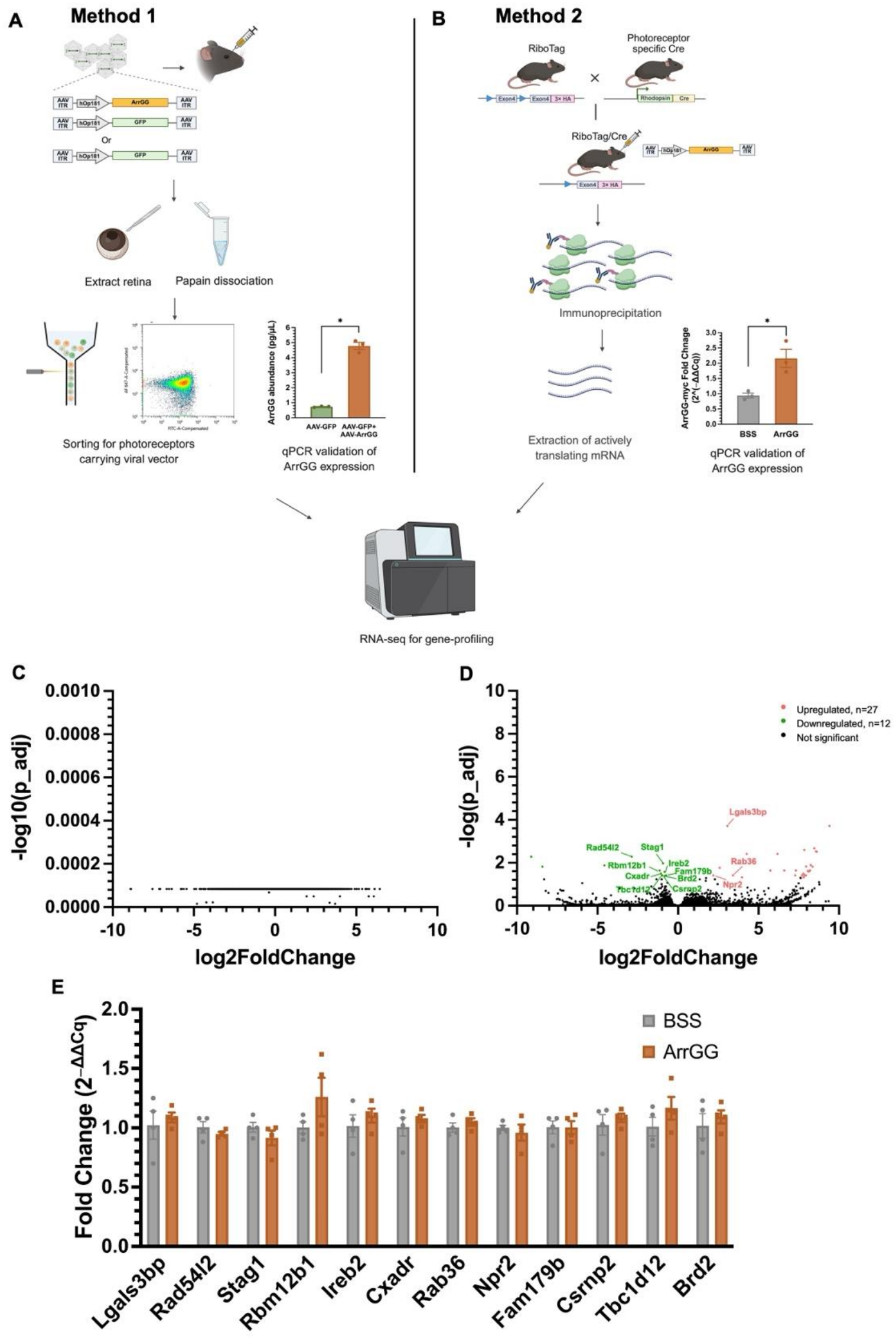
ArrGG does not induce major transcriptomic changes in photoreceptors of C57BL/6J mice. **(A)** Method 1: Schematic of the FACS-based workflow for photoreceptor RNA isolation. AAV-ArrGG was co-injected with AAV-GFP into the vitreous of C57BL/6J mice. GFP-positive/CD73-positive photoreceptors were isolated by FACS for bulk RNA sequencing. ArrGG expression was verified by RT-qPCR. Transcriptomic profiles were compared between the ArrGG/GFP-treated and GFP-treated groups. Data are presented as mean ± SEM. Statistical significance was determined by the unpaired *t*-test (*: *P*<0.05). **(B)** Method 2: Schematic of the RiboTag-based workflow for photoreceptor mRNA isolation. RiboTag mice were intravitreally injected with AAV-ArrGG and ribosome-associated photoreceptor mRNAs were isolated for bulk RNA sequencing. ArrGG expression in RiboTag-enriched photoreceptor mRNA was confirmed by RT-qPCR. Transcriptomic profiles were compared with those from RiboTag mice injected intravitreally with balanced salt solution (BSS). Data are presented as mean ± SEM. Statistical significance was determined by the unpaired *t*-test (*: *P*<0.05). **(C)** Volcano plot of RNA-seq analysis from Method 1 comparing the ArrGG/GFP-treated group (*n*=3) with the GFP-treated group (*n*=3). **(D)** Volcano plot of RNA-seq analysis from Method 2 comparing the ArrGG-treated group (*n*=3) with the BSS-treated group (*n*=3). Differentially expressed genes were defined as |log₂FC| ≥ 1 and adjusted *P* ≤ 0.05. **(E)** RT-qPCR analysis of candidate genes identified in panel **D** using independent biological samples (*n*=4). Data are presented as mean ± SEM. Statistical significance was determined by multiple unpaired *t*-tests.

Since this approach identified no differentially expressed genes, which is unusual for a transcriptomic analysis, we chose to validate the absence of transcriptional effects of ArrGG using a complementary photoreceptor RNA isolation method based on the RiboTag system^31^. Because photoreceptor mRNA was isolated using the RiboTag strategy rather than GFP-based cell sorting, mice treated with balanced salt solution (BSS) were used as the control group. The RiboTag mouse carries a hemagglutinin (HA)-tagged *Rpl22* allele that enables isolation of ribosome-associated mRNA from specific cell types^18^. For our experiments, RiboTag mice were crossed with rhodopsin-Cre mice^32^ to express HA-tagged ribosomes predominantly in rod photoreceptors, allowing photoreceptor mRNA to be isolated by HA immunoprecipitation (**Fig. 2B**). Four weeks after intravitreal injection of AAV-ArrGG (or BSS), ribosome-associated photoreceptor mRNA was isolated from rhodopsin-Cre/*Rpl22* mice for transcriptomic analysis. ArrGG expression was confirmed in RiboTag-isolated photoreceptor mRNA by RT-qPCR analysis, with significantly higher ArrGG expression detected in ArrGG-treated samples compared with the BSS-treated controls (**Fig. 2B**).

Using this complementary approach, the overall differential expression analysis showed that there were very few genes meeting the predefined thresholds for differential expression in ArrGG-treated samples compared with BSS-treated controls (**Fig. 2D**). To more carefully validate if any of the genes identified in Fig. 2D were differentially expressed in ArrGG-treated eyes, we performed RT-qPCR analysis of the candidate genes using a separate set of samples isolated through the same RiboTag procedure. However, the RT-qPCR analysis did not confirm the significant expression changes observed by RNA-seq (**Fig. 2E**), indicating that the candidate transcriptional changes identified by RNA-seq were not reproducible.

Together, these two complementary transcriptomic approaches revealed no evidence of widespread transcriptional remodeling in the photoreceptor populations following ArrGG expression and are consistent with the proposed idea that the principal effect of ArrGG is on metabolic activity rather than transcriptional remodeling.

### ArrGG Preserves Retinal Function and Slows Photoreceptor Loss in an RPE-Based IRD Model

Because ArrGG targets metabolic enhancement, we have proposed that ArrGG might offer a gene-agnostic approach for slowing retinal degeneration by improving the metabolic environment of both photoreceptors and the surrounding Müller glia and RPE cells. We previously showed that ArrGG expression slowed retinal degeneration in the *Rho^P23H/+^* mouse model^17^. Since rhodopsin is the cognate binding partner for arrestin1, it is possible that the effect we observed in this particular model of IRD might be related to this natural interaction, rather than to the effect on glycolytic output. To determine whether the effects of ArrGG extend to other genetic forms of IRD, we delivered ArrGG to the retinas of three genetically distinct mouse models of retinal degeneration using the same AAV construct.

We first evaluated the therapeutic efficacy of ArrGG in the retinal degeneration 6 (rd6) mouse. The rd6 mouse carries a splice donor mutation in the membrane-type frizzled-related protein (*Mfrp*) gene, which is predominantly expressed in the retinal pigment epithelium^33, 34^. Consequently, rd6 serves as a model of retinitis pigmentosa arising from an RPE defect and is characterized by slow, progressive photoreceptor degeneration^35^. In this mouse, retinal dysfunction is detectable by one month of age and progresses to complete loss of photoreceptor outer segments by fifteen months of age^35^.

For this study, rd6 mice were injected at P30 before substantial photoreceptor loss had occurred and were followed until P330, when photoreceptor degeneration was readily evident. Mice received unilateral intravitreal injections of AAV-ArrGG, AAV-ArrWT, or BSS, while the contralateral eye remained untreated as an internal control.

Representative dark-adapted ERG traces recorded at P210 showed that ArrGG-treated rd6 mouse eyes exhibited larger response amplitudes than the contralateral untreated eyes, as well as the BSS-treated and ArrWT-treated eyes (**Fig. 3A**). Intensity-response functions showed similar results, with ArrGG-treated eyes exhibiting greater a-wave and b-wave amplitudes than untreated, BSS-treated, and ArrWT-treated eyes, particularly at higher flash intensities (**Fig. 3B, C**). Dark-adapted ERG responses at 30 cd·s/m² were monitored longitudinally to evaluate whether the functional benefit of ArrGG was sustained (**Fig. 3D, E**). Eyes that received ArrGG showed a slower decline in both a-wave and b-wave amplitudes compared with the untreated control eyes throughout the study period. This slowing of ERG decline was not observed in eyes that were treated with either BSS or ArrWT (**Fig. 3D, E**), supporting a treatment-associated benefit of ArrGG. Although ArrGG-treated rd6 eyes exhibited greater ERG amplitudes than untreated eyes, their responses remained below those of the wild-type C57BL/6J mice included as a reference for normal retinal function.

**Figure 3.**
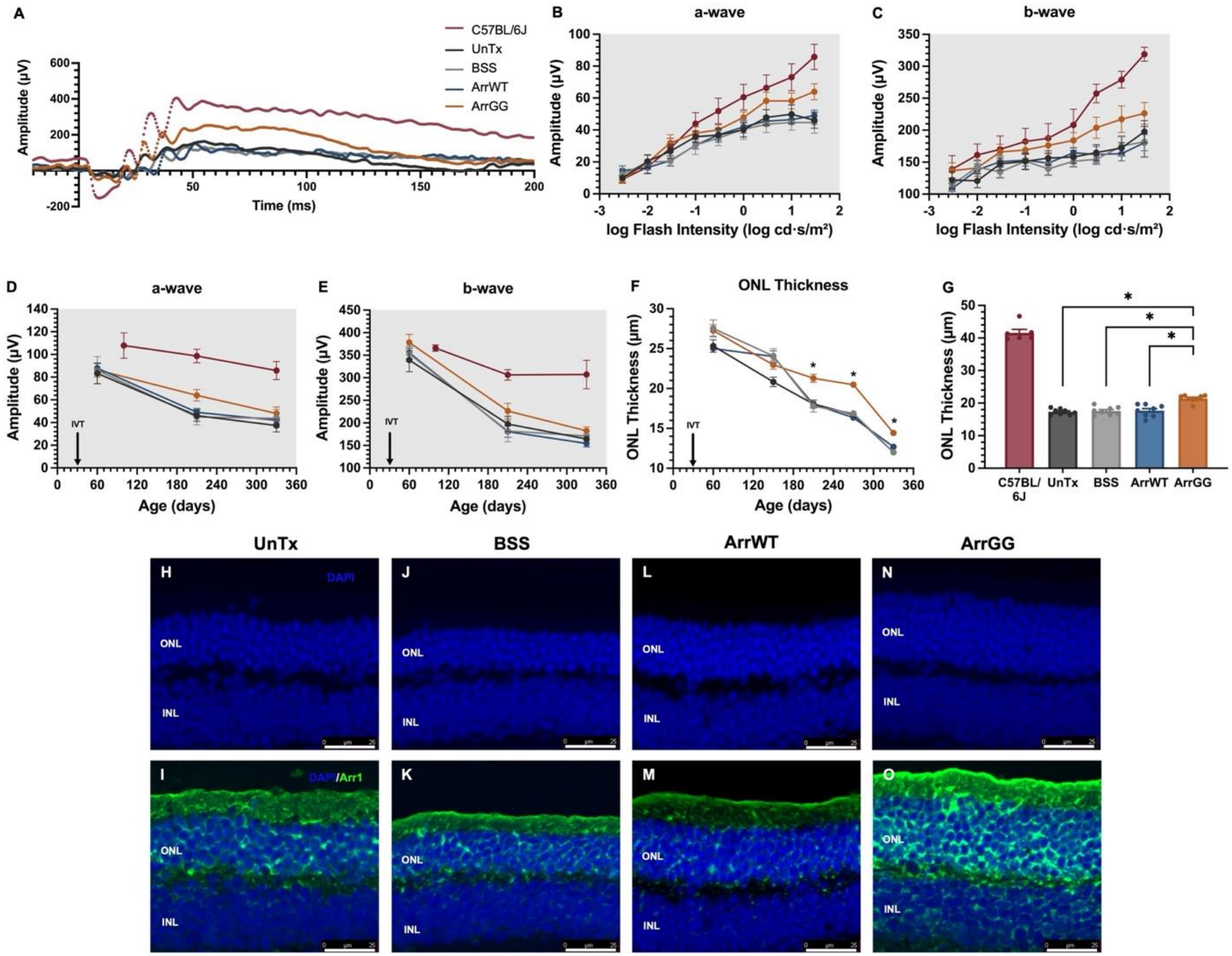
ArrGG slows retinal degeneration in *Mfrp^rd6^*^/J^ (rd6) mice. **(A**) Representative dark-adapted ERG responses recorded from untreated wild-type C57BL/6J mouse eyes (P330) and untreated (UnTx), BSS-treated, ArrWT-treated, and ArrGG-treated rd6 mouse eyes (P210), with treatments administered at P30. **(B, C)** Intensity-response functions for dark-adapted ERG a-wave (**B**) and b-wave (**C**) amplitudes recorded from wild-type C57BL/6J mouse eyes (P330, *n*=8) and untreated, BSS-treated, ArrWT-treated, and ArrGG-treated rd6 mouse eyes (P210, *n*=16). **(D, E)** Longitudinal measurements of dark-adapted ERG a-wave **(D)** and b-wave **(E)** amplitudes at 30 cd·s/m^2^ in wild-type C57BL/6J mouse eyes (P100, P210, and P330, *n*=8) and untreated, BSS-treated, ArrWT-treated, and ArrGG-treated rd6 mouse eyes (P60, *n*=13; P210, *n*=16; P330, *n*=10). **(F)** Longitudinal measurements of outer nuclear layer (ONL) thickness by OCT at approximately two-month intervals in untreated, BSS-treated, ArrWT-treated, and ArrGG-treated rd6 mouse eyes (*n*=8). ONL thickness was quantified at 1,400 μm temporal to the optic nerve head. Data are presented as mean ± SEM. Statistical significance was determined by two-way ANOVA (*: *P*<0.05). (**G**) ONL thickness measured by OCT in wild-type C57BL/6J mice (P180, *n*=6) and untreated, BSS-treated, ArrWT-treated, and ArrGG-treated rd6 mouse eyes (P210, *n*=8). Data are presented as mean ± SEM. Statistical significance was determined by one-way ANOVA (*: *P*<0.05). Wild-type C57BL/6J data are shown for reference only and were excluded from statistical analyses. **(H-O)** Representative immunohistochemical images of retinal sections collected at P210 from untreated **(H, I)**, BSS-treated **(J, K)**, ArrWT-treated **(L, M)**, and ArrGG-treated **(N, O)** rd6 mouse eyes. Sections were stained with DAPI (blue) and anti-arrestin1 (green).

Because the preservation of retinal function is expected to correlate with reduced photoreceptor degeneration, we also monitored ONL thickness by OCT at approximately two-month intervals (**Fig. 3F**). Retinas in both control groups (BSS-treated and ArrWT-treated) showed a continuous decline in ONL thickness, consistent with the previous report^35^. Although ONL thinning was also observed in ArrGG-treated eyes, the rate of thinning was slower than that in the control groups. A significantly thicker ONL became evident from P210 onward in ArrGG-treated eyes compared with untreated controls (**Fig. 3F**); however, ONL thickness in ArrGG-treated rd6 eyes remained below the wild-type C57BL/6J reference values (**Fig. 3G**). Together, the ERG and ONL thickness measurements indicate that ArrGG slows retinal degeneration in the RPE-based rd6 model by preserving retinal function and attenuating photoreceptor loss, without fully preventing retinal degeneration.

Immunohistochemical analysis of retinal sections from rd6 mice showed findings consistent with our OCT measurements (**Fig. 3H-O**). At P210, ArrGG-treated retinas retained approximately eight rows of photoreceptor nuclei within the ONL (**Fig. 3N, O**), whereas untreated, BSS-treated, and ArrWT-treated retinas contained only four to six rows (**Fig. 3H-M**), consistent with the greater ONL thickness observed by OCT. Together, these findings demonstrate that ArrGG slows retinal degeneration in rd6 mice by preserving retinal function and attenuating photoreceptor loss.

### ArrGG Slows Photoreceptor Loss in a Cone-Associated IRD Model

To further investigate whether the therapeutic benefit of ArrGG extends across genetically distinct forms of inherited retinal degeneration, we next evaluated the efficacy of ArrGG in the guanylate cyclase-1 knockout (GC1KO) mouse model. The GC1KO mouse carries a null mutation in the *Gucy2e* gene, resulting in progressive cone dysfunction and degeneration, and serves as a model of cone-associated early-onset Leber congenital amaurosis type 1 (LCA1)^36–38^. As in the previous model, AAV-ArrGG was injected intravitreally into one eye, with the contralateral eye left untreated as an internal control, and compared with eyes receiving AAV-ArrWT or BSS. Treatment was administered at P14 before significant cone photoreceptor loss, and mice were followed until P126.

Although the GC1KO mouse does not show evidence of rod photoreceptor loss because of the presence of GC2 in rods^37^, we measured dark-adapted intensity-response functions in our cohort of ArrGG-treated mice to confirm this expectation in our colony of GC1KO mice. No significant differences in dark-adapted amplitudes were observed among the treatment groups (**Fig. 4A**). However, dark-adapted b-wave amplitudes in GC1KO mice remained below the wild-type C57BL/6J reference (**Fig. 4A**). This observation is consistent with previous reports showing that rod function is retained but reduced in the GC1KO mouse model^37^.

**Figure 4.**
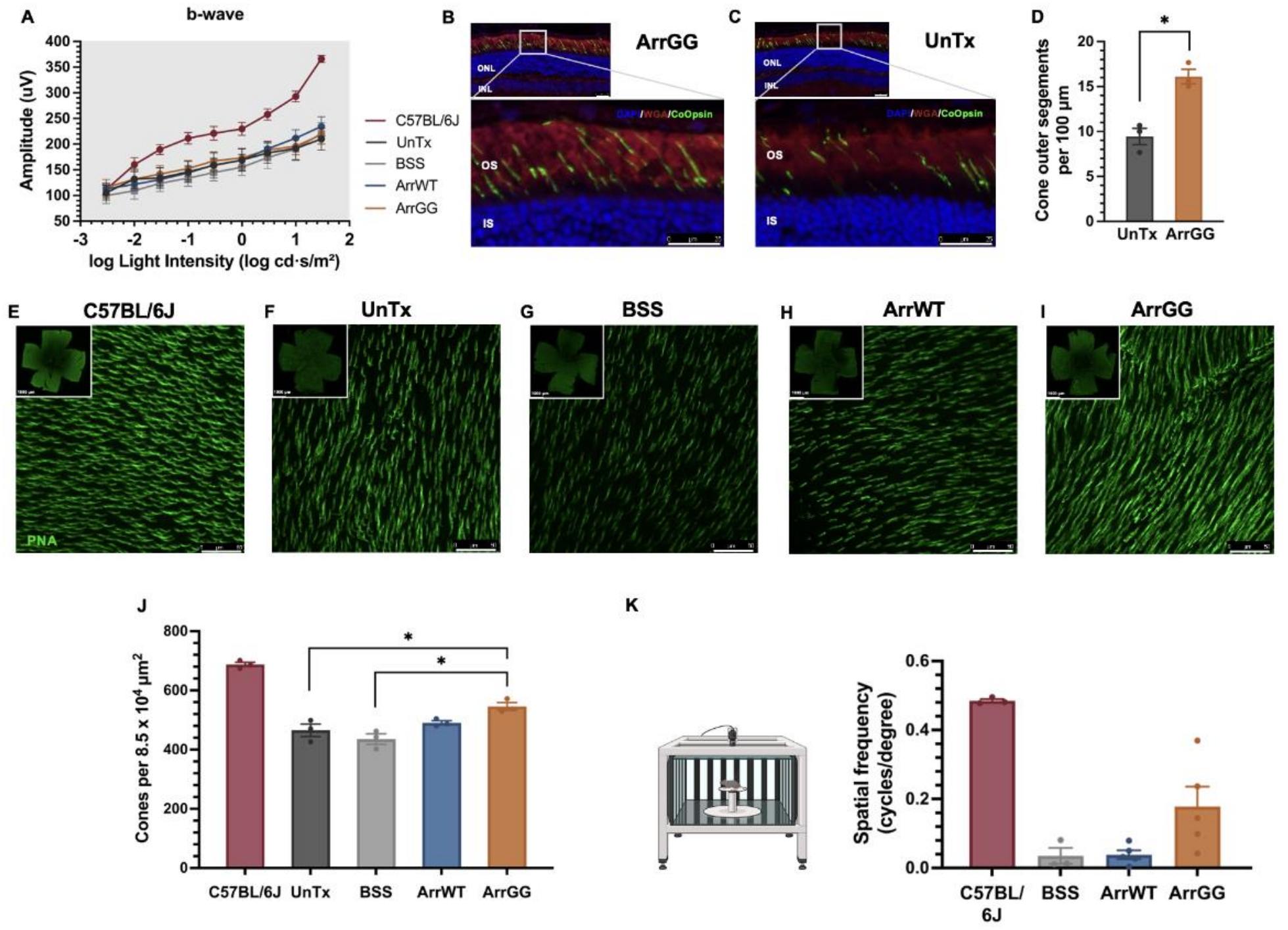
ArrGG slows retinal degeneration in guanylate cyclase-1 knockout (GC1KO) mice. **(A)** Intensity-response functions for dark-adapted ERG b-wave amplitudes recorded from wild-type C57BL/6J mouse eyes (P100, *n*=10) and untreated, BSS-treated, ArrWT-treated, and ArrGG-treated GC1KO mouse eyes (P98, *n*=10), with treatments administered at P14. **(B, C)** Representative immunohistochemical images of retinal sections collected at P126 from ArrGG-treated **(B)** and untreated **(C)** GC1KO mouse eyes. Sections were stained with anti-cone opsin (green), DAPI (blue), and WGA (red). (**D**) Quantification of cone outer segments from immunohistochemical images of retinal sections (n=3). Data are presented as mean ± SEM. Statistical significance was determined by the unpaired *t*-test (*: *P*<0.05). **(E-I)** Representative retinal flat mounts stained with PNA (green) from wild-type C57BL/6J mouse eyes at P50 **(E)** and untreated **(F)**, BSS-treated **(G)**, ArrWT-treated **(H)**, and ArrGG-treated **(I)** GC1KO mouse eyes at P98. **(J)** Cone numbers were quantified based on PNA-positive staining in the flat mounts shown in panels **E-I** (*n*=3). Data are presented as mean ± SEM. Statistical significance among GC1KO groups was determined by one-way ANOVA (*: *P*<0.05). **(K)** Schematic of the optomotor response testing system used to assess visual acuity in mice (left). Spatial frequency thresholds (right) were measured in BSS-treated (*n*=3), ArrWT-treated (*n*=5), and ArrGG-treated (*n*=5) GC1KO mice at P98. Wild-type C57BL/6J mice (*n*=3) were included as a reference and were excluded from statistical analyses.

In contrast, GC1KO mice exhibit progressive cone photoreceptor loss^36^. To assess whether ArrGG could promote cone photoreceptor preservation in the absence of GC1, we examined cone morphology using immunohistochemical methods. Retinal sections collected at P126 were stained with anti-cone opsin to visualize cone outer segments (**Fig. 4B, C**). Representative retinal sections showed substantially more cone opsin-positive outer segments in ArrGG-treated eyes than in untreated eyes, with approximately 16 cone outer segments per 100 μm of retinal length in ArrGG-treated retinas, compared with approximately 9 per 100 μm in contralateral untreated retinas (**Fig. 4D**).

Because cones are not uniformly distributed throughout the mouse retina, we also prepared retinal flat mounts to assess cone preservation across a broader retinal area. Flat mounts were collected from GC1KO mice at P98 and stained with peanut agglutinin (PNA), which selectively binds to the cone-specific interphotoreceptor matrix (IPM)^39^ (**Fig. 4E-I**). Wild-type retinal flat mounts were included as a reference for normal cone distribution and density. The number of PNA-positive cone structures was significantly higher in ArrGG-treated eyes than in untreated and BSS-treated eyes (**Fig. 4J**), though the counts remained below the wild-type reference values. Although PNA labeling does not directly measure cone integrity, it is widely used as an indicator of cone preservation, and these findings were consistent with our cone opsin immunohistochemistry results.

Finally, to determine whether the preservation of cone photoreceptors was accompanied by improved visually guided behavior, GC1KO mice were evaluated using an optomotor response test. At P98, GC1KO mice exhibited markedly reduced spatial frequency thresholds relative to the wild-type reference values. Although mice receiving ArrGG had lower spatial frequency thresholds than wild-type C57BL/6J mice, the optomotor response in the ArrGG group was improved compared to GC1KO mice receiving either BSS or ArrWT (**Fig. 4K**).

Together, these findings demonstrate that ArrGG promotes cone photoreceptor preservation and improves visual performance in the GC1KO mouse model.

### ArrGG Preserves Retinal Function and Slows Photoreceptor Loss in a Severe Rod Degeneration Model

To evaluate the therapeutic efficacy of ArrGG in a severe model of retinitis pigmentosa, we next delivered ArrGG to the retinas of the retinal degeneration 10 (rd10) mouse model. The rd10 mouse carries a spontaneous mutation in the rod phosphodiesterase 6β (*Pde6b*) gene, leading to rapid rod degeneration beginning around P18, followed by secondary cone loss and retinal remodeling^40^.

For therapeutic evaluation in this rapidly degenerating rod disease model, ArrGG was delivered intravitreally at P15 before structural degeneration became evident, with the contralateral eye left untreated as an internal control. Control mice received intravitreal injections of BSS. Mice were followed until P206, when advanced retinal remodeling was evident^41^.

At P195, dark-adapted a-wave amplitudes in all rd10 groups remained at or near noise level and did not increase with flash intensity (**Fig. 5A**). However, b-wave amplitudes were significantly greater in ArrGG-treated eyes at the two highest flash intensities, 10 cd·s/m² and 30 cd·s/m², which elicit a mixed rod-cone response (**Fig. 5B**). Light-adapted ERG measurements also revealed greater b-wave amplitudes in ArrGG-treated eyes at higher flash intensities. Taken together, these findings suggest a preservation of cone-mediated retinal function (**Fig. 5C**). Wild-type C57BL/6J mice exhibited substantially greater dark-adapted and light-adapted ERG amplitudes than rd10 mice and were included as a reference to illustrate the extent of functional impairment associated with retinal degeneration in rd10 mice.

**Figure 5.**
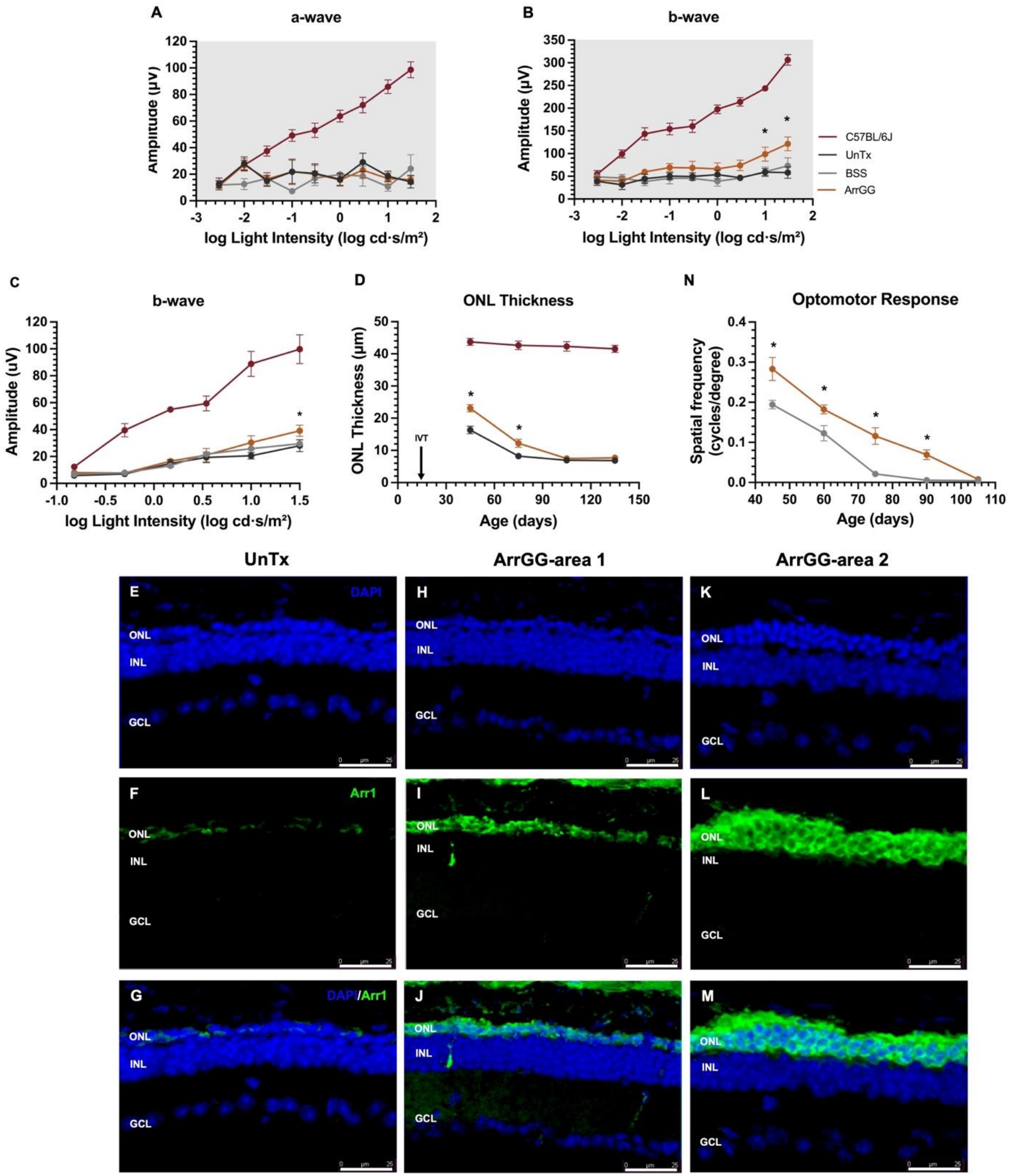
ArrGG slows retinal degeneration in *Pde6b^rd^*^10/J^ (rd10) mice. **(A, B)** Intensity-response functions for dark-adapted ERG a-wave **(A)** and b-wave **(B)** amplitudes recorded from wild-type C57BL/6J mouse eyes (P210, *n*=10) and untreated, BSS-treated, and ArrGG-treated rd10 mouse eyes (P195, *n*=6). (**C**) Intensity-response functions for light-adapted ERG b-wave amplitudes recorded from wild-type C57BL/6J mouse eyes (P150, *n*=10) and untreated, BSS-treated, and ArrGG-treated rd10 mouse eyes (P75, *n*=6). **(D)** Longitudinal measurements of ONL thickness by OCT at approximately monthly intervals in wild-type C57BL/6J mouse eyes (P45, *n*=6; P75, *n*=8; P105, *n*=6; P135, *n*=6) and untreated and ArrGG-treated rd10 mouse eyes (P45, *n*=15; P75, *n*=9; P105, *n*=7; P135, *n*=9). ONL thickness was quantified at 1,200 μm temporal to the optic nerve head. Data are presented as mean ± SEM. Statistical significance was determined by two-way ANOVA (*: *P*<0.05). Wild-type C57BL/6J data are shown for reference only and were excluded from statistical analyses. **(E-M)** Representative immunohistochemical images of retinal sections collected at P206 from untreated rd10 mouse eyes **(E-G)** and ArrGG-treated rd10 mouse eyes (**H-M**). In eyes treated with ArrGG, most of the retina showed modest photoreceptor preservation (**H-J**), although there were some smaller regions noted that showed more significant photoreceptor survival (**K-M**). Sections were stained with DAPI (blue) and anti-arrestin1 (green). (**N**) Longitudinal measurements of optomotor spatial frequency thresholds in BSS-treated and ArrGG-treated rd10 mice (P45, P60, P75, P90, P105; *n*=5 per group). Data are presented as mean ± SEM. Statistical significance was determined by two-way ANOVA (*: *P*<0.05).

ONL thickness was monitored by OCT at monthly intervals (**Fig. 5D**). Because rod degeneration in rd10 begins around P18 and peaks near P25^40^, substantial photoreceptor loss had already occurred at P45 when we performed our first OCT measurement. Despite this advanced stage of degeneration, ArrGG-treated eyes retained a significantly thicker ONL than untreated eyes (**Fig. 5D**). This preservation was maintained through P105, consistent with slower progression of photoreceptor loss in ArrGG-treated rd10 mice. Although ONL thickness remained substantially below that of wild-type C57BL/6J mice, ArrGG treatment provided measurable structural preservation during disease progression.

To further characterize photoreceptor preservation at an advanced stage of degeneration, retinal sections collected at P206 were examined by immunohistochemistry (**Fig. 5E-M**). In our examination, we noted that untreated eyes generally contained a single row of nuclei in the ONL with minimal detectable arrestin1 staining (**Fig. 5E-G**). In contrast, eyes treated with ArrGG exhibited varying degrees of photoreceptor preservation. Regions with limited preservation retained approximately 1-2 rows of ONL nuclei (**Fig. 5H-J**), whereas regions with stronger preservation and more prominent arrestin1 staining contained approximately 3-4 rows of ONL nuclei (**Fig. 5K-M**). These small areas with more retained photoreceptors did not show any particular topographic distribution, but instead were scattered across the retina. These preserved regions exhibited more prominent arrestin1 staining, consistent with the retention of photoreceptor cells. The variation in local ONL preservation indicates that the structural benefit of ArrGG treatment was not uniform across the retina.

To determine whether the observed retinal preservation was accompanied by sustained visually guided behavior, longitudinal optomotor measurements were performed from P45 to P105 (**Fig. 5N**). ArrGG-treated rd10 mice exhibited higher spatial frequency thresholds than BSS-treated mice during the progression of retinal degeneration, with measurable responses persisting to later ages. However, optomotor thresholds progressively declined in both groups, approaching minimal levels at advanced stages of degeneration. It should be noted that although ArrGG-treated rd10 mice retained measurably higher dark-adapted b-wave amplitudes at P195 (**Fig. 5B**), the absence of an improved optomotor response at P105 suggests that the preservation of retinal electrophysiological responses did not necessarily correspond to sustained improvement in visual function at least as measured by optomotor responses.

Together, these findings demonstrate that ArrGG preserves photoreceptor-mediated retinal function, slows photoreceptor loss, and delays the decline in optomotor responses in the rapidly degenerating rd10 mouse model.

## Discussion

Our previous work has established a model in which the glycolytic potential of photoreceptors is partially reduced by an interaction between arrestin1 and enolase1^15–17^. Because of the central role of glycolysis in the metabolism of the retina, we proposed that disinhibiting enolase1 with a modified arrestin1 (ArrGG) could be used as a method to increase glycolysis for therapeutic benefit. In this study, we extend our previous findings to show that the effect of ArrGG is directly on glycolysis using a cell culture system. We also show that ArrGG can be used across multiple models of IRD to slow the rate of photoreceptor loss and the decline of visual function. This effect is most likely attributable to the increased glycolytic output of photoreceptors treated with ArrGG, as significant transcriptional changes were not associated with ArrGG application.

The cellular experiments provide some mechanistic insight into the effect of ArrGG on metabolism. Our current study shows that the addition of arrestin1 to ARPE-19 cells reduces lactate production in these cells, consistent with our prior studies using HEK-293T cells^16^. This current study further demonstrates that this effect of arrestin1 is due to a reduction in the rate of glycolysis. Furthermore, the addition of ArrGG to these suppressed cells completely restored the compensatory glycolytic response. Obviously, ARPE-19 cells do not adequately reflect the complexity of a layered retina, so it is important to not over-interpret these findings. Together with the previously established interaction between ArrGG and enolase1^16^, these findings are consistent with the hypothesis that ArrGG counteracts ArrWT-mediated inhibition of enolase1 activity, likely through competition for enolase1 binding.

An important remaining question is the downstream fate of glucose-derived carbon following ArrGG-associated modulation of glycolysis. This is particularly relevant in the retina because photoreceptors, Müller glia, and retinal pigment epithelial cells form a closely integrated metabolic ecosystem^10, 11, 42^, such that changes in photoreceptor glycolysis may influence multiple metabolic pathways and neighboring retinal cells. Isotope tracing using labeled glucose^10, 43^ will therefore be important for examining carbon flow following ArrGG treatment and assessing its broader impact on retinal metabolism. Photoreceptors rely heavily on aerobic glycolysis not only for energy production but also for supporting biosynthesis^44^, redox homeostasis^7^, and continuous outer segment renewal^44^. Therefore, future metabolomics studies coupled with stable isotope tracing will allow direct quantification of glucose-derived carbon utilization following ArrGG treatment and provide a more comprehensive understanding of how altered glycolytic metabolism contributes to retinal homeostasis and photoreceptor survival.

In addition to these mechanistic findings, we also explored if ArrGG expression could have a broader influence since metabolic adaptation is frequently associated with alterations in the expression of genes involved in glycolytic and non-glycolytic pathways, including mitochondrial metabolism, cellular stress responses, and epigenetic regulation^45–48^. Using two complementary and independent RNA-seq approaches, we showed that the application of ArrGG to the retina does not lead to widespread transcriptomic alterations in photoreceptors. Although the RiboTag approach identified a few candidate genes that might be changed in expression level, changes in the expression of these genes were not confirmed by the more rigorous RT-qPCR analyses and are thus considered as false positives. Regardless, the consistency of the results between these two orthogonal approaches—one identifying no changes and the other identifying very few changes—provides confidence in concluding that the application of ArrGG does not cause significant transcriptional changes in photoreceptors.

It is important to note that our transcriptomic analyses specifically focused on photoreceptor-enriched RNA to evaluate potential changes in the primary cell population targeted by ArrGG since ArrGG expression was mainly targeted to photoreceptors using an opsin promoter. Although whole-retina RNA-seq could identify additional transcriptional responses, changes detected in other retinal cell types might reflect secondary effects of altered photoreceptor metabolism rather than direct effects of ArrGG expression. Such indirect responses may contribute to retinal protection and remain an important area for future investigation. During the preparation of this manuscript, we also noted a recent publication reporting unintended Cre recombination in approximately 6% of RPE cells in the rhodopsin-iCre mouse line used in our RiboTag model ^49^. It is our assessment that this limited off-target activity is unlikely to substantially affect the overall interpretation of our transcriptomic analysis, given the consistent findings obtained using an independent FACS-based photoreceptor enrichment approach. Overall, these observations are consistent with the hypothesis that the effects of ArrGG do not require widespread changes in photoreceptor gene expression, although they do not establish the underlying mechanism of retinal protection.

One of the important findings in this current study is that ArrGG application resulted in the slowing of retinal degeneration in three different models of IRD. The protective effects of ArrGG observed in the rd6 model are an important finding since the primary genetic defect in this mouse model originates outside photoreceptors. In the rd6 model, *Mfrp* deficiency is associated with the loss of ADIPOR1 protein from the apical membrane of RPE cells and progressive photoreceptor degeneration^50–52^. Enhancing glycolysis directly within photoreceptors may increase their metabolic resilience and ability to tolerate conditions associated with RPE dysfunction. Rather than correcting the underlying RPE defect, ArrGG may promote photoreceptor survival by supporting metabolic processes required for cellular maintenance under the chronic stress imposed by RPE dysfunction.

Likewise, ArrGG provided protection to cones in the GC1KO mouse model, in which cones undergo selective degeneration while rods remain largely preserved due to the presence of a second retinal guanylate cyclase, GC2^36^. Our findings demonstrated greater cone preservation following ArrGG treatment along with improved visual function as assessed by optomotor responses. In our analysis of the optomotor response, we noted that the thresholds measured in our control GC1KO mice were lower than those previously reported by Boye et al.^53^. Although the reason for this discrepancy remains uncertain, differences in animal cohorts, observer-dependent response assessment, or other experimental factors may have contributed, despite the use of consistent procedures and blinded assessment in the present study. Regardless of the reason for this disparity, our findings indicate that ArrGG provided protective effects.

The protective effects of ArrGG may reflect the metabolic interactions within the photoreceptor microenvironment. Although glucose represents a major metabolic substrate for photoreceptors, cones can also utilize lactate as a metabolic fuel^54^. Thus, enhanced glycolytic activity and increased lactate production following ArrGG treatment may improve local metabolic substrate availability and contribute to cone survival. In addition to arrestin4 (a.k.a. cone arrestin), arrestin1 is also abundantly expressed in cones^55^, suggesting that direct modulation of the arrestin1-enolase1 interaction in cones could also have therapeutic potential. We do not anticipate that substantial disinhibition of enolase1 occurs directly in cones with the current ArrGG vector, because the human opsin promoter drives ArrGG expression predominantly in rod photoreceptors. Together, these observations suggest that ArrGG-mediated modulation likely supports cone survival through changes in the local metabolic environment, extending the potential application of ArrGG to models of cone degeneration.

Finally, this current study also showed that ArrGG could prolong photoreceptor survival and slow vision loss in a model of rapid photoreceptor degeneration. The rd10 mouse carries a mutation in *Pde6b* that disrupts rod PDE6 activity, leading to abnormal cGMP and Ca^2+^ homeostasis and rapid rod degeneration followed by secondary cone loss^40, 56^. In contrast to the relatively slow progression of photoreceptor loss in *Rho^P23H/+^*mice or the rd6 mouse, the rapid degeneration in rd10 mice provides a substantially narrower therapeutic window. The preservation of photoreceptors and their function following ArrGG treatment therefore suggests that the benefit of metabolic modulation is not restricted to slowly progressing diseases. Consistent with this concept, Zhang et al. demonstrated that increasing glycolytic flux promoted photoreceptor survival in a PDE6-associated retinal degeneration model^57^. Interestingly, treatment-associated improvements in ERG responses were most apparent in the b-wave at higher flash intensities, whereas a-wave amplitudes did not show a significant treatment effect. The greater b-wave amplitudes at higher flash intensities may reflect contributions from surviving cones and residual rod-mediated signaling. Since cone degeneration occurs secondary to rod loss in rd10 mice, it is also possible that slowing rod degeneration with ArrGG supports cone survival and function. However, the present experiments do not establish the relative contributions of rods and cones. Furthermore, although histological examination revealed greater ONL preservation in ArrGG-treated retinas, this preservation was regionally variable, with areas of limited and more pronounced photoreceptor survival, which may not produce a substantial retinal response that can be recorded by full-field ERG measurements. Spatially resolved functional measurements would be required to establish a direct relationship between regional structural preservation and local retinal function.

Together, this study extends our previous work by demonstrating treatment-associated structural and functional preservation across genetically distinct mouse models of inherited retinal disease, supporting the potential of ArrGG as a gene-agnostic therapeutic approach. It is important to note that although our study shows structural and functional benefits associated with ArrGG, this modified arrestin1 does not stop retinal degeneration but rather slows the rate of photoreceptor loss. Further metabolic flux analysis will be necessary to clarify the metabolic consequences of ArrGG expression and determine the extent to which these changes contribute to retinal protection.

